# Mechanical advantage is not invoked by additional cognitive load during multi-finger dexterous object manipulation

**DOI:** 10.1101/2025.10.21.683756

**Authors:** Swarnab Dutta, Prajwal Shenoy, SKM Varadhan

**Author notes:** Corresponding author; Prajwal Shenoy **Email**. **Author Contributions:** Conceptualisation – SD, VSKM; Methodology – SD, VSKM; Investigation – SD, PS; Formal Analyses – SD, PS; Original Draft – SD, PS; Review and Editing – SD, PS, VSKM. **Competing Interest Statement:** The authors declare no competing interests.

## Abstract

Everyday interactions with objects depend on keeping the grasped item in static equilibrium by effectively distributing the contact forces from the fingertips. The mechanical advantage hypothesis (MAH) proposes that the central nervous system (CNS) optimises the distribution of grasping forces by favouring fingers with longer moment arms. While existing research has established that the applicability of MAH is influenced by biomechanical constraints, it remains unclear whether cognitive load, a distinct dimension of task difficulty, can similarly induce this force-sharing strategy. To investigate this, we examined whether increased cognitive demands during a multi-finger grasp-and-lift task would trigger force distribution patterns that exploit mechanical advantage. The results showed that while participants completed the task successfully, contrary to our hypothesis, the increased cognitive load did not result in a disproportionate rise in the normal force exerted by the little finger compared to the ring finger. This finding suggests that the employment of optimisation strategies like mechanical advantage may be predominantly sensitive to biomechanical rather than cognitive constraints.

**Significance Statement:** Human grasping is a redundant control problem in that the hand has more ways to produce forces and moments than are strictly needed to achieve desired object interaction. The mechanical advantage hypothesis (MAH) is an optimisation principle proposing that the central nervous system (CNS) prefers fingers with greater leverage to address this redundancy during object manipulation. While MAH is known to emerge when mechanical difficulty increases, this study probes its boundaries and generality by asking whether nonmechanical demands also trigger it. The findings show that increased cognitive demand does not invoke MAH, highlighting that this optimisation arises primarily in response to physical and biomechanical task demands, with implications for dexterous control in rehabilitation, assistive technology, and robotics.

## Introduction

The human hand’s remarkable capacity for grasping and manipulating objects is fundamental to our daily interactions. Such dexterity depends on the precise coordination of fingertip forces to achieve and sustain static equilibrium while handling objects, ensuring stable and efficient manipulation. This allows us to engage in a range of activities, from delicate precision tasks to powerful grips. Achieving such a stable grasp involves a dynamic interplay of sensorimotor processes, enabling the rapid and flexible rearrangement of fingertip forces in response to both expected and unpredictable perturbations. Research on object manipulation has examined the individual forces contributed by the fingers and thumb when varying the properties of the grasped objects. These studies have systematically altered grasp conditions through changes in object orientation[1], [2], mass[3], [4], mass distribution [5], [6], [7], [8], surface friction [9], [10], [11], grip aperture [12], individual digit width [13], external torque[8], [14] and finger or thumb positions[15], [16]. Such perturbations often induced instability, necessitating compensatory moments from the fingers to rebalance the manipulandum. A key conceptual framework for understanding grasping with a redundant system like the human hand in producing moments on an object is the mechanical advantage hypothesis (MAH) [17], [18]. The MAH proposes that the central nervous system (CNS) coordinates finger forces during the production of a moment in such a way that fingers with longer moment arms exert greater normal forces than those with shorter moment arms. Specifically, when the thumb functions as a pivot, the peripheral fingers, such as the index and little fingers, have longer moment arms for exerting normal force compared to the central fingers, which are the middle and ring fingers. The mechanical advantage hypothesis (MAH) asserts that the little finger enhances its effectiveness by increasing its normal force during movements that require supination, particularly in comparison to the adjacent ring finger. This adjustment enables the ulnar fingers to exert less overall normal force while still maintaining the necessary supination moment for achieving static equilibrium. MAH thus suggests an optimised force distribution that facilitates efficient moment production while maintaining object stability with minimal overall effort.

To explore the boundaries and generality of MAH, several studies have investigated its applicability in five-finger grasping tasks. Support was observed in experiments involving the rotation of a manipulandum in both pronation and supination directions at two different speeds.[19]. Additionally, studies also explored MAH through the application of external loads at different distances from the centre of mass [14] and producing moments to follow a trapezoidal template by applying force with all four fingers in a pressing task [20]. However, the hypothesis only received partial support in tasks involving moment production with a mechanically fixed object[21], where the distance from the fingers to the axis of rotation was systematically altered, alongside changes in the magnitude and direction of moment production. The partial or conditional applicability of MAH in this study led the authors to propose its task-specific and effector-dependent nature. Recent experimental studies have further refined our understanding of when and why mechanical advantage is invoked. For instance, a previous study from our group [16] observed that a clear manifestation of mechanical advantage (where the little finger produces more normal force than the ring finger) emerged only at the highest tested load (0.450kg). At lower loads (0.150kg, 0.250kg, 0.350kg), the force contributions from the ulnar fingers were comparable, suggesting that the CNS engages mechanical advantage only past a threshold of mechanical difficulty. In a follow-up study, [22] Rajakumar et al. demonstrated that imposing a constraint wherein the thumb normal force was limited to a nominal value invoked MAH even at the moderate load (0.250kg), supporting the idea of a threshold specification, since such task constraints can raise the effective mechanical demands to, or beyond, the required threshold. These results collectively suggest that the CNS flexibly deploys the mechanical advantage strategy in response to heightened task difficulty.

While task difficulty in determining the applicability of MAH has been explored in relation to mechanical factors in previous studies, examining another dimension through the role of cognitive load, which refers to additional attentional or mental demands placed on the individual, may offer valuable insights. In everyday life, we often handle objects while multitasking, such as conversing, remembering instructions, or responding to environmental distractions. It is plausible that increased cognitive demand would elevate the effective task difficulty and, in turn, would shape how the CNS organises grasp forces to maintain stability. A growing body of literature documents the disruptive impact of cognitive load on upper limb performance, including reaching and grasping[23], [24], [25], [26], [27], precision and power grip squeezing [28], force tracking [29], [30], [31], [32], and tapping [33], [34], [35]. Specifically, grip-lift experiments conducted alongside concurrent working memory tasks show that cognitive load can prolong task phases and elicit greater grip forces, even during skilled, familiar interactions [36], [37], [38]. These findings suggest that higher-order cognition interacts dynamically with sensorimotor processes, indicating that grasp control is not fully automatic but sensitive to the allocation of cognitive resources.

Given that cognitive loading alters grip kinetics and movement timing, it remains an open question whether such loading can also affect how forces are distributed among digits during multi-finger grasping. Specifically, we do not yet know if cognitive demands can induce the CNS to switch to a mechanical advantage like strategy at moderate loads where it would not typically be employed. If so, this would reveal a previously unrecognised pathway via cognitive and not only biomechanical constraints, by which the CNS optimises grasp force distribution. The present study thus investigates whether additional cognitive load at a moderate mechanical load (0.250 kg) is sufficient to invoke this strategy during multi-finger grasping. We hypothesise that cognitive demand would push the effective task difficulty, leading to preferential recruitment of the little finger (longer moment arm) relative to the ring finger (shorter moment arm), thereby revealing mechanical advantage like force sharing even when mechanical constraints alone would not elicit such a response.

## Results

### Handle Orientation and Thumb Platform Displacement

The results of the tilt calculation show that the net tilt was well below 2 degrees for all cases, as shown in Table 1, which indicates that the handle was maintained in static equilibrium. The thumb displacement values also indicate that the thumb sliding platform was maintained well within the limit throughout all the different conditions. These indicate that the participants successfully performed the motor task in all three experimental conditions.

**Table 1.** Average and S.E.M. values for the net tilt and thumb platform displacement error for all three conditions. S.E.M refers to the Standard Error of Means.

| Cond | Net Tilt (degrees) |  | Thumb displacement error (cm) |  |
| --- | --- | --- | --- | --- |
|  | Average | S.E.M | Average | S.E.M |
| ST | 0.379 | 0.216 | 0.253 | 0.028 |
| DTL | 0.854 | 0.267 | 0.258 | 0.029 |
| DTH | 0.720 | 0.277 | 0.299 | 0.025 |

### Analysis of grip forces

In order to investigate the effect of cognitive load on grip force throughout the task, four separate one-way repeated-measures ANOVAs were conducted on virtual finger grip force (VF_GF) at lift onset, lift off, maximum grip force, and during the hold phase with Condition [Control (ST), Low Cognitive Load (DTL), High Cognitive Load(DTH)] as the within-subject factor. For lift onset, there was no significant effect of Condition (F (2,34) =0.53, p=0.59, η_p_^2^=0.03). At lift off, the effect of Condition was not significant (F (2,34) =0.31, p=0.73, η_p_^2^=0.01). For maximum grip force, Condition did not reach significance (F (2,34) =0.13, p=0.88, η_p_^2^=0.01), and during the hold phase, results were similarly non-significant (F (2,34) =0.31, p=0.73, η_p_^2^=0.01). Thus, none of these analyses revealed any significant effects of Condition, and all effect sizes (partial eta squared) were small, indicating stable VF grip force across cognitive load levels at each key epoch. Equivalence tests were performed to rigorously examine the virtual finger grip force (VF_GF) at four key task epochs across all paired condition comparisons (Control vs. Low Load, Control vs. High Load, Low Load vs. High Load). For virtual finger grip force at load force onset, the comparison between Control and Low Load yielded t (17) = -2.26, p < 0.05, Control versus High Load showed t (17) = − 2.01, p < 0.05, and Low Load compared to High Load resulted in t(17)=-2.79,p<0.01. At lift off, Control and Low Load had t(17)=2.46,p<0.05; Control and High Load showed t(17)=2.25,p<0.05; and Low Load versus High Load yielded t(17)=2.62,p<0.01. For maximum grip force, Control versus Low Load was t(17)=2.45,p<0.05, Control versus High Load was t(17)=1.85,p<0.05, and Low Load versus High Load was t(17)=2.682,p<0.01. During the hold phase, Control and Low Load produced t(17)=2.71,p<0.01, Control and High Load produced t(17)=2.51,p<0.05, and Low Load versus High Load showed t(17)=2.66,p<0.01. All tests confirmed equivalence at the p < 0.05 level, supporting similar virtual finger grip force across cognitive load manipulations at each task phase.

To further investigate the main hypothesis, a Two-way repeated-measures ANOVA was conducted on grip force with Finger (Ring, Little) and Condition [Control (ST), Low Cognitive Load (DTL), High Cognitive Load(DTH)] as within-subject factors. The two-way repeated-measures ANOVA was conducted with Finger (ring, little) and Condition (control, low, high difficulty) as within-subjects factors. There was no significant main effect of Finger (F(1,17)=0.78,p=0.39,η_p_2=0.02), and the main effect of Condition was not significant (F(2,34)=1.57,p=0.22,η_p_2=0.04. The interaction between Finger and Condition also did not reach significance (F(2,34)=0.76,p=0.39,η_p_2=0.02. These results indicate that neither finger type nor experimental condition, nor their interaction, produced significant differences in the outcome variable across the three levels tested. For Ring and Little finger comparisons in each condition, the ST comparison revealed t(17)=-2.00,p<0.05, the DTL comparison yielded t(17)=−1.74,p<0.05, and the DTH comparison t(17)=2.53,p<0.05. All tests confirmed equivalence at the p < 0.05 level, supporting comparable ulnar finger forces for all three experimental conditions.

## Discussion

The present study examined whether heightened cognitive demands at a moderate load (0.250 kg) would elicit a shift in force-sharing strategies consistent with the mechanical advantage hypothesis (MAH). It was reasoned that cognitive load, by imposing additional attentional costs, can be conceived as another dimension of elevating task difficulty apart from strictly mechanical loading. Overall, the results demonstrated that participants were able to perform the grasping task successfully under cognitive interference, i.e., without loss of stability or failure in achieving equilibrium (Table 1). Thus, the central nervous system (CNS) appeared capable of flexibly integrating additional cognitive demands while maintaining successful task performance.

With respect to immediate task performance, one notable outcome was the absence of an overall increase in grip forces under cognitive load. This absence of increase in grip force was observed consistently across all four epochs considered for evaluating total grip force (Fig. 3), specifically the virtual finger grip force at lift onset, lift off, maximum grip force, and the central hold phase. This finding contrasts with prior grip–lift studies reporting higher grip force magnitudes when participants simultaneously engaged in secondary cognitive tasks. We speculate that this discrepancy may be attributable to the mechanical characteristics of our manipulandum. Specifically, a free-to-move thumb-slider configuration was employed, which, as reported in [39], already induces greater grip forces compared to a fixed-slider design. This setup was used to deliberately amplify the contribution of the ulnar fingers in compensatory moment production, as our focus was to check for mechanical advantage within the ulnar fingers. To elaborate, constraining the thumb slider restricted the tangential force contribution by the thumb, thereby rendering it relatively inconsequential for producing the supination (clockwise) moment. This design, in turn, amplified the necessity for greater ulnar fingers’ grip force engagement in producing the compensatory moment. Consequently, because participants were already operating at a relatively elevated level of grip force, there may have been little scope left for further increases when a cognitive load was imposed. Put differently, the additional cognitive demands may have created a ceiling effect, masking the incremental changes in grip-force magnitudes that are more typically observed in dual-task studies. It should be noted that participants began the motor task with a substantial grip force already applied to the object, as the experimental protocol did not include a distinct reach phase (Fig. 5)

Turning to the main hypothesis, additional cognitive load did not invoke a mechanical advantage– like strategy at moderate load (Fig. 4). In other words, contrary to our expectations, the little finger did not generate a statistically significant greater normal force compared to the ring finger under increased cognitive demand. Why though? One possibility is that the elevated baseline forces associated with the free-slider manipulandum obscured more subtle effects, thereby reducing the detectability of shifts in relative force recruitment between the ulnar digits. Future experiments could help address this by investigating this in the fixed-slider condition, where lower baseline grip-force requirements might create sufficient scope for detecting alterations in digit-force organisation under cognitive load.

An alternative interpretation, which we think is more plausible, is that optimisation strategies such as MAH may be preferentially recruited under conditions of purely biomechanical challenge, as opposed to cognitive constraints. Existing evidence suggests that the mechanical advantage principle emerges robustly when mechanical difficulty is sufficiently high, for instance, at heavy external loads, or when explicit limits are placed on thumb force contribution [22], [40]. By contrast, cognitive load may not by itself constitute a sufficient trigger for invoking an optimisation strategy by the CNS. From this perspective, we suggest that the underlying computation of force-sharing is primarily tuned to biomechanical requirements of stability and efficiency, while cognitive interference only modulates temporal measures or overall force magnitudes [36], [37], [38]. This rationale thus implies that while the CNS accommodates cognitive load, it does so through conservative stabilisation strategies, rather than by reorganising digit forces to exploit biomechanical advantage. Taken together, these findings raise the possibility that optimisation strategies such as MAH are not equally sensitive to all forms of task difficulty. Rather, they may be more selectively deployed in response to physical and biomechanical constraints, such as heavier loads or explicit force limitations. In contrast, when the additional difficulty arises from cognitive sources, the CNS may instead prioritise robustness and stability over fine-tuned optimisation of force distribution. In conclusion, the present study suggests that increasing cognitive demands at moderate load does not alter the organisation of digit forces according to the principle of mechanical advantage. Instead, optimisation strategies in multi-finger grasping may be preferentially reserved for conditions where mechanical demands surpass a critical threshold.

Several limitations warrant mention, which we plan to address in our forthcoming research. The use of the free-to-move slider manipulandum may have contributed to a ceiling effect in grip force, potentially obscuring subtle digit-level reorganisation that could reveal mechanical advantage under heightened cognitive demands. Replicating the experiment with a fixed-slider condition, especially at lower baseline grip forces, may yield clearer insights. Another possibility is that the cognitive load manipulation in the present study may not have been sufficiently challenging to impose significant demands on executive resources, which raises the need to test the paradigm under higher or more complex cognitive loads. Future studies could incorporate pupillometry to quantify cognitive load and strategy use more directly. Additionally, exploring a wider range of mechanical loads and levels of cognitive interference would help determine the precise threshold at which the mechanical advantage strategy is recruited. Finally, given that the present study did not directly measure neural activity or actual motor-unit activation patterns, further research integrating neurophysiological markers is required to clarify the mechanisms through which mechanical advantage emerges. Future investigations should also aim to disentangle the relative influences of cognitive and biomechanical constraints by systematically manipulating both within the same experimental paradigm, and across manipulandum designs that vary in baseline grip-force requirements. This approach would address whether distinct gateways, vis-à-vis cognitive and biomechanical, exist for invoking mechanical advantage or whether the strategy is fundamentally biomechanical in nature.

## Materials and Methods

### Participants

Eighteen right-handed participants (12 males and 6 females, age ± S.D. = 25.6 ± 3.8) took part in this study. Written informed consent was obtained from all participants prior to the start of the study. None of the participants reported any history of neuromotor disorders or injuries to the arm or hand. Handedness was assessed using the Edinburgh Handedness Inventory[41]. The experimental procedure was approved by the Institutional Human Ethics Committee (IITM-IHEC) at the Indian Institute of Technology Madras (Approval Number: IEC/2023-01/SKM/20). All experimental sessions were conducted in accordance with the guidelines established by the Institute’s Human Ethics Committee. Participants received financial compensation in accordance with the Institute’s guidelines for their involvement in a study that lasted approximately one hour.

### Experimental setup

The experiment employed a custom-designed five-finger precision manipulandum constructed from aluminium, as shown in Fig. 1. The manipulandum was equipped with five force/torque sensors (Nano 17, ATI Industrial Automation, North Carolina, USA) that measured forces and moments along the X, Y, and Z axes with a resolution of 0.0125 N for both normal and tangential components. The thumb sensor was mounted on a sliding platform via a baseplate, whereas the sensors for the other four fingers were directly fixed to the handle using a single baseplate. To measure the vertical displacement of the thumb platform, a laser displacement sensor (Baumer, India; Model OADM 12U6460) with 5 μm resolution was positioned on top of the manipulandum. A bull’s-eye spirit level was integrated into the handle’s upper section, enabling participants to monitor and maintain rotational alignment. Handle orientation was recorded using a BNO055 IMU (Bosch Sensortec; 16-bit, 2000°/s range). With an additional load of 250 g, the total weight of the handle was 765 g. Data from the force/torque sensors, displacement sensor, and IMU were synchronised and acquired at 100 Hz using a custom LabVIEW program.

**Figure 1:**
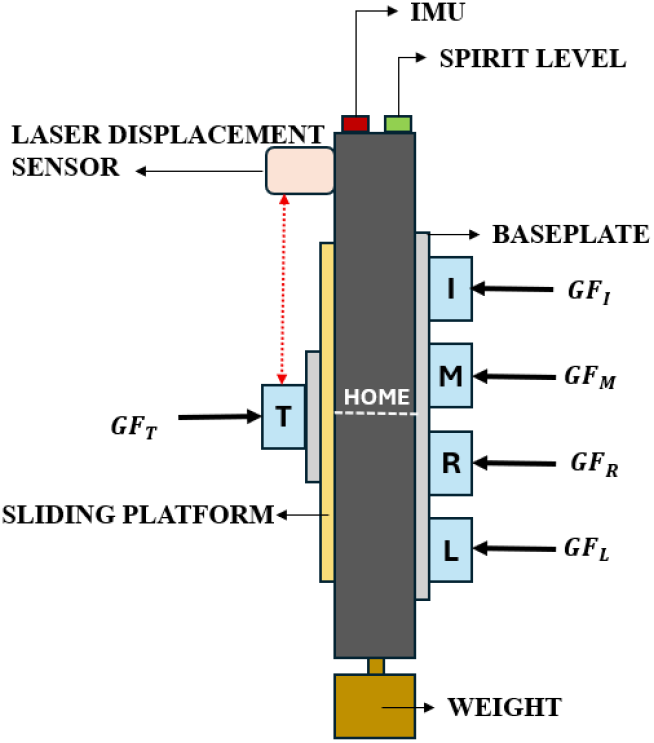
Schematic image of the five-fingered grasping handle equipped with force sensors for each finger: Thumb -T, Index-I, Middle-M, Ring-R, and Little-L. The thumb sensor was mounted on a sliding platform. A laser displacement sensor was utilised to measure the movement of the slider. An Inertial Measurement Unit (IMU) measured the tilt of the handle. Additionally, a spirit level was included to help participants maintain the handle in the vertical position before starting the motor task. The grip forces acting on each finger and thumb are shown in the figure to illustrate their line of action.

### Experimental Design

Each experimental session of a participant was split into 3 conditions, viz., the control condition, Single-task ST, the low difficulty condition, named Dual-task low DTL, and the high difficulty condition, named Dual-task high DTH. Each condition involved 12 trials. The ST condition involved only a motor task, whereas the DTL and DTH conditions involved a motor task with low and high cognitive loading, respectively, that loaded the verbal working memory. In all three conditions, the motor task involved grasping the manipulandum and lifting it. The participants were initially instructed to sit in a relaxed manner on a wooden chair with their hands placed on the table. Participants were then instructed to grasp a five-fingered manipulandum using the fingertips of their thumb and four fingers, and lift the manipulandum to a comfortable height in a natural and relaxed manner, while doing their best to prevent it from tilting in any direction. For the cognitive loading, the string recall task from the literature [38] was adapted. In this task, the participants were required to memorise a sequence of letters before the beginning of the motor task. In the high-difficulty condition, a series of consonant letters was displayed in red on a grey background in a dynamic manner. The sequence was constructed from the chosen letters J, K, L, M, N, P, Q, R, S, T, and V. To ensure a higher level of difficulty and engagement, vowels and familiar acronyms were omitted, preventing participants from using mnemonic techniques that involve creating pronounceable chunks or pseudo-words. The task involving LOW cognitive loading was constructed using the same letters but in their correct alphabetical order, for instance, K, L, M, N. This allowed the participants to rely on their semantic knowledge for memorising the displayed sequence. The length of the sequence for both the DTL and DTH conditions was tailored to match each participant’s assessed memory span, which was determined prior to the session and remained constant throughout the experiment. The sequence of the three conditions ST, DTL and DTH was balanced across all participants. A 5-minute break was provided after each condition.

In the ST condition, the participants initially grasped the manipulandum with the thumb platform aligned to the centre, in preparation for the trial. Upon holding the manipulandum, the participants were directed to fix their gaze on a red fixation cross on the screen. After hearing the first auditory cue, the participants lifted the handle, held it in the air, and finally, upon hearing the second auditory cue, they lowered and released the handle in its initial lowered position. The time between the 2 auditory cues was 6 seconds. The forces and moments from the 5 sensors, along with the tilt of the handle (measured using the IMU) and the thumb displacement (measured using the laser displacement sensor), were recorded during the 6-second window.

In the DTL and DTH conditions, the participants initially grasped the manipulandum with the thumb platform aligned to the centre, preparing for the experiment. Upon holding the handle, the participants were directed to fix their gaze on a red fixation cross on the screen. After the participants confirmed having positioned their fingertips on the sensors, a sequence of text was displayed for 2.25 seconds. The participants were required to memorise the sequence. Upon hearing an auditory cue, they lifted the handle, held it in the air, and upon hearing the second auditory cue, they lowered and released the handle. After completing the motor task, a single letter from the presented sequence was displayed on the screen for 1 second and then made to disappear. The participants were required to verbally recite the letter that followed the displayed letter in the original sequence. For the motor task, if the tilt exceeded a particular limit or if the thumb slider slid beyond the threshold limit, the trial was discarded and repeated. For the cognitive task, the inability to recall the correct letter resulted in the rejection and repetition of the trial. The experimental protocols for the ST, DTL and DTH experimental conditions are shown in Fig. 2.

**Figure 2:**
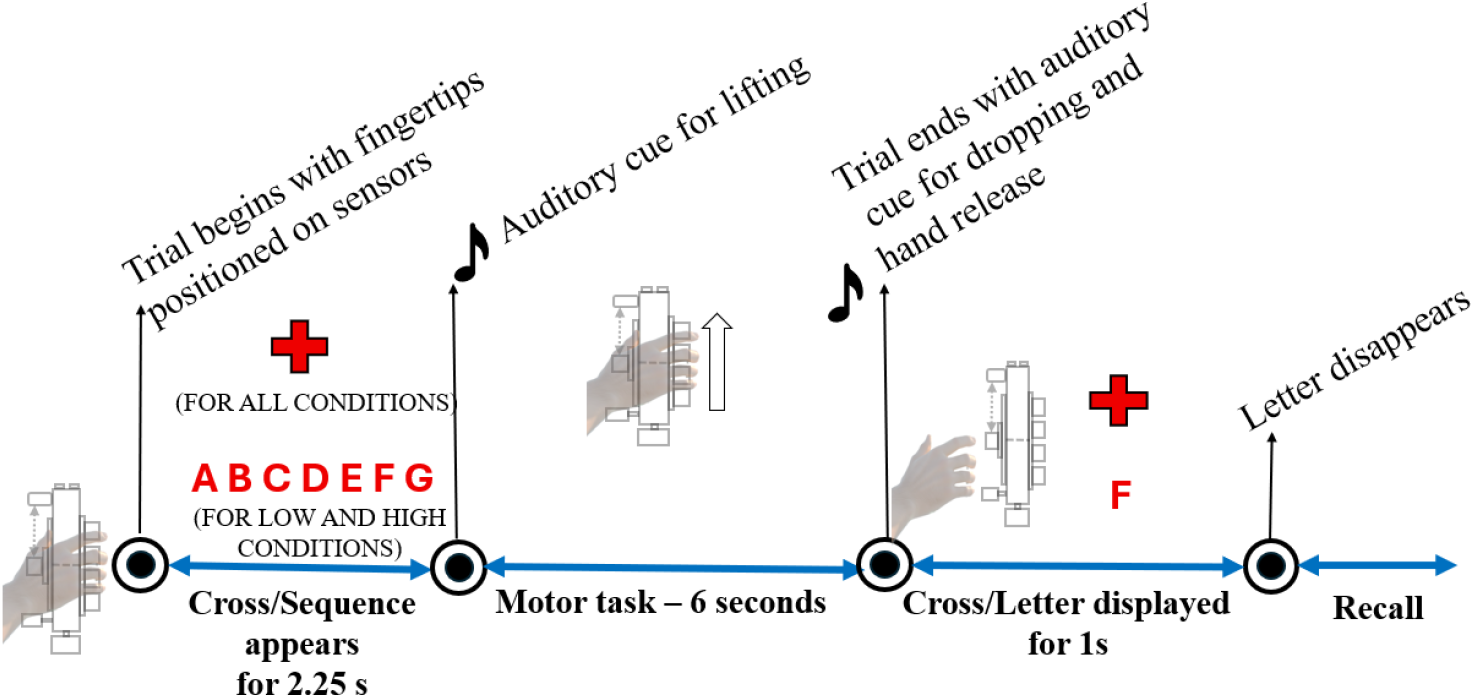
Experimental protocol used in the study. For the ST task involving only a motor task, the participants held the handle and shifted their gaze to a fixation cross on the screen. Upon hearing an auditory cue, they lifted the handle and then released it upon hearing the second auditory cue. For the tasks involving the motor task as well as cognitive loading (Dual-task low DTL and Dual-task high DTH), after the participants positioned their fingers on the sensors, a sequence of letters appeared for 2.25 seconds and then disappeared. The participants, after memorising the sequence, performed the motor task for 6 seconds, following which a letter from the sequence was displayed, and the recall test was performed.

**Figure 3:**
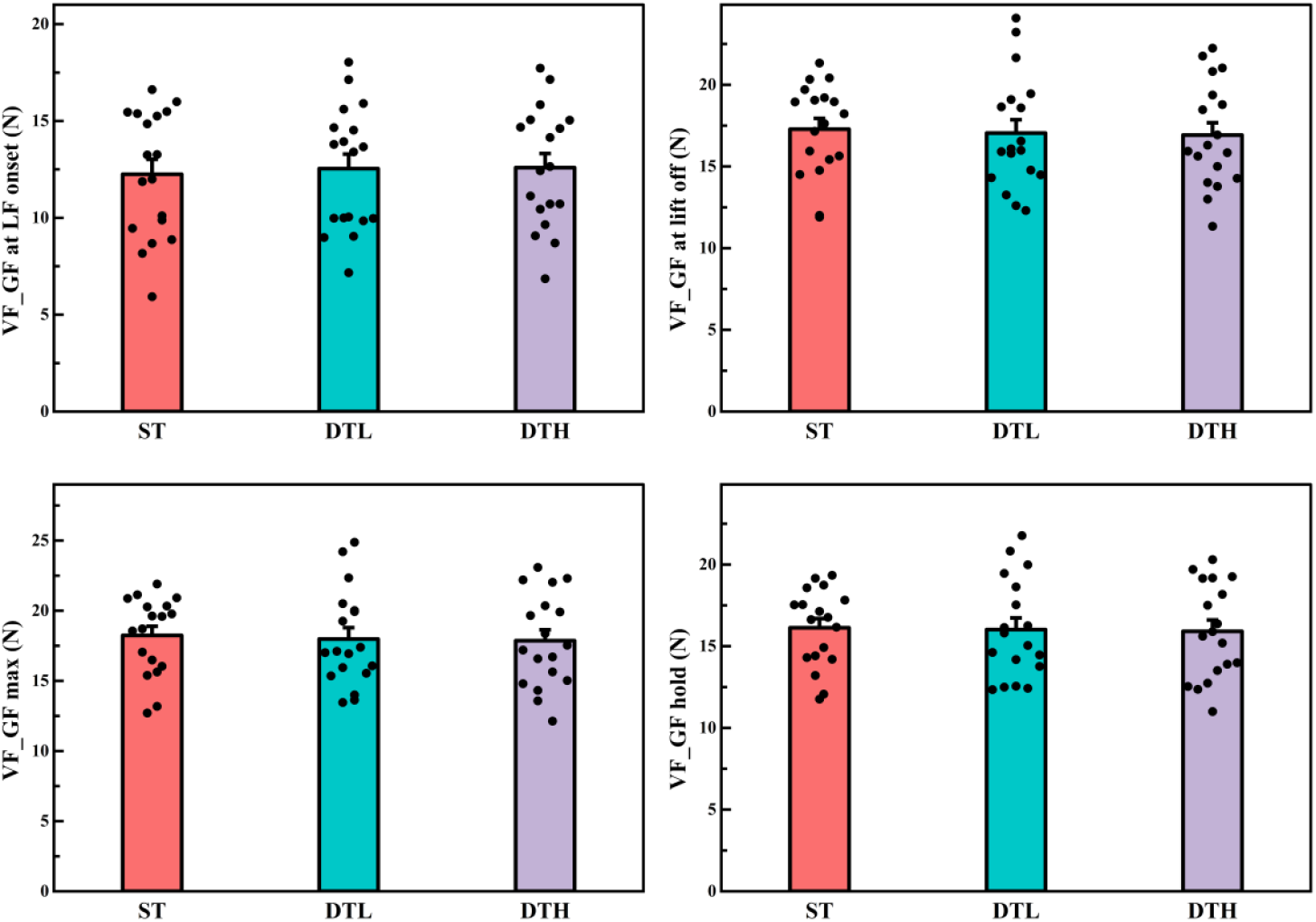
Average of the virtual finger grip force at load force onset (LF onset), at lift off, during hold phase, and maximum virtual finger grip force during the trial across the three experimental conditions. The vertical columns and bars indicate the means and the standard errors of means (SE), respectively. Note that the horizontal jitter applied to the data points is solely a visualisation technique to prevent overlap and does not represent additional experimental data.

**Figure 4:**
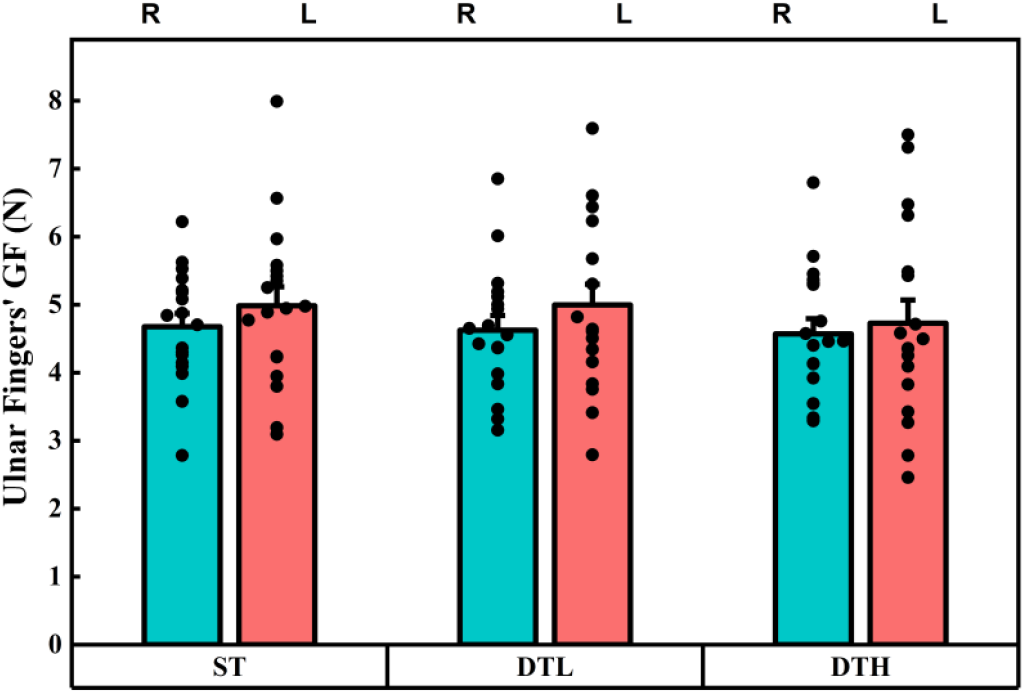
Average of the ulnar fingers’ grip forces during the hold phase across the three experimental conditions. The vertical columns and bars indicate the means and the standard errors of means (SE), respectively. Note that the horizontal jitter applied to the data points is solely a visualisation technique to prevent overlap and does not represent additional experimental data.

**Figure 5:**
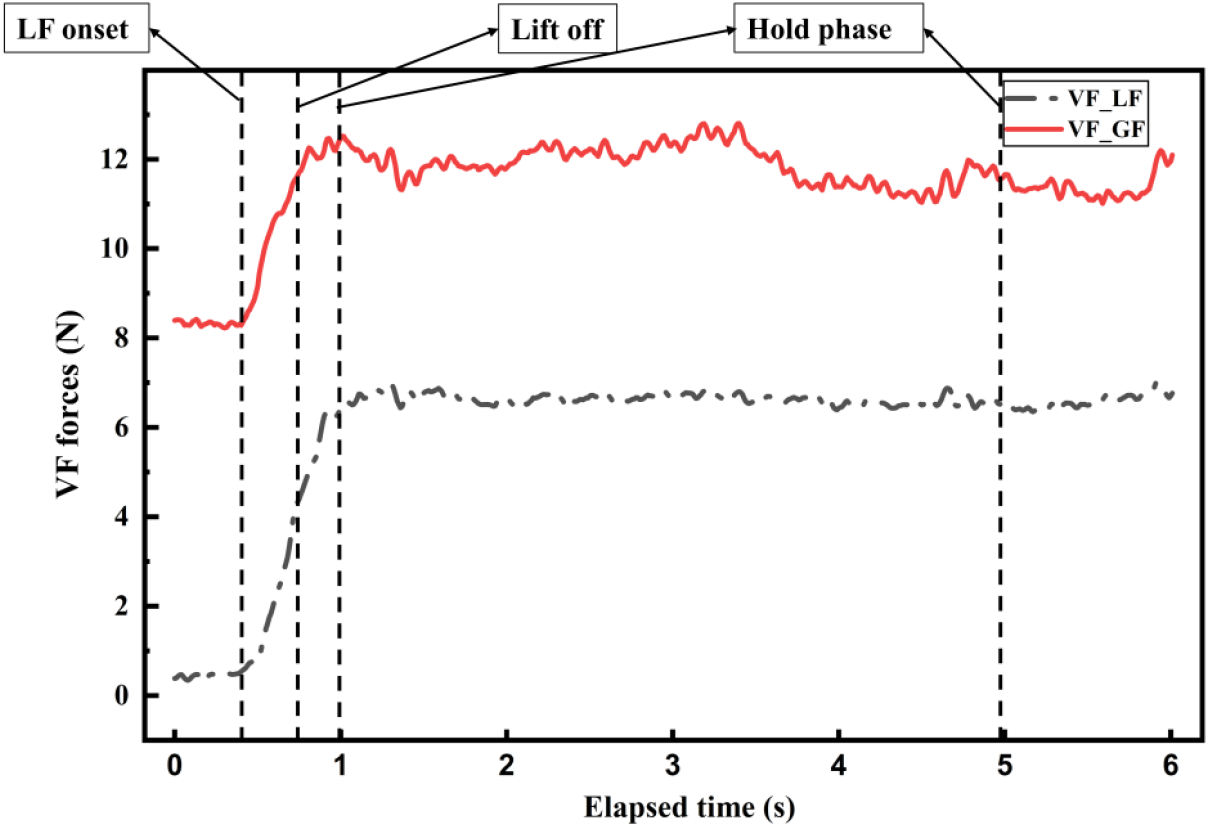
Traces of the virtual finger grip force (solid line) and load force (dashed line) during a representative trial from the dual task high difficulty condition. Vertical dotted lines indicate crucial epochs of the experiment, including load force onset, lift off, and hold phase.

### Pre-Experimental Assessment of Memory Span

Before the beginning of the experiment, each participant was assessed for memory span using the Mini-Mental State Examination (MMSE) [42]. For this purpose, they were required to memorise a text sequence and reproduce verbally the memorised sequence. The procedure initially began with displaying a sequence with three letters for 2.25 seconds. The participants recited it after the sequence disappeared and a fixation cross appeared on the screen. Every time the participant successfully recited the memorised sequence, the length of the sequence was increased by one. If the participant failed to memorise and verbally reproduce the sequence of length ‘n’, the assessment was stopped, and the previous length of the sequence (n-1) was considered as the memory span for that participant. Before the experiment, the cognitive function of all participants was assessed using the MMSE screening, in addition to the memory span evaluation.

### Data Analysis

The signals from the force/torque sensors and the laser displacement sensor were filtered using a second-order low-pass Butterworth filter with a 15 Hz cutoff and zero-phase lag. An Inertial Measurement Unit (IMU) was set to output quaternions, which were used to determine the orientation of the handle. Before starting the experiment, the handle was positioned at zero degrees along the X, Y, and Z axes for a duration of 5 seconds, with alignment verified visually using a spirit level. The quaternions obtained from this calibration period were averaged to define the reference quaternion. During the experimental trials, the recorded orientation quaternions were expressed relative to this reference using the quaternion conjugate operation.

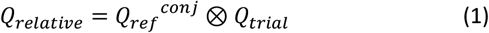

The handle’s net tilt was calculated using relative quaternions. Trials were discarded and repeated if the net tilt exceeded 3 degrees. For each participant, the net tilt during the central 4 seconds of the motor task was averaged, and these averages across participants were compiled for all conditions. The results are presented as average net tilt values along with the standard error of the mean to evaluate the handle’s stability. Similarly, thumb displacement values were averaged and reported. Any thumb displacement that deviated by ± 0.5 cm from the midline position (6 cm) led to the rejection of that trial. In the analysis, only the grip forces of the ring, little, and thumb fingers were taken into account. For each condition, the central 4 seconds of the 6-second motor task were designated as the hold phase to eliminate effects from the beginning and end of the trials. The data was averaged for each condition individually and then aggregated across participants. The mean values and standard error of the mean are shown for comparison. To evaluate the mechanical advantage hypothesis, grip forces from the little and ring fingers were compared across all conditions and presented in a single graph.

The analysis of grip and load forces during the manipulation task was segmented into distinct epochs defined based on the time course of force events. The epochs were defined as Load Force Onset: This event was defined as the time point when the load force (LF) of the virtual finger exceeded the baseline mean plus four standard deviations (mean + 4 SD). The baseline period was established from a pre-movement window where forces were stable, ensuring load onset reflects a significant force increase, indicating object lift initiation. Lift Off: Lift off was defined as the instant when the total load force, calculated as the sum of the thumb and virtual finger load forces, surpassed the object’s weight. Maximum Grip Force: The peak grip force generated by the virtual finger during the hold and lift phases was identified and analysed as an index of the force scaling to maintain object stability. Central Hold Phase: This phase corresponds to the time window from 1 second to 5 seconds, representing a steady-state period where grip and load forces stabilise during object holding. The virtual finger is a conceptual framework used to simplify the analysis of multi-digit grasping. Instead of analysing each finger individually, the forces from the four fingers, i.e., index, middle, ring, and little, are combined vectorially to form a single “virtual finger” [43]. This allows the calculation of collective grip and load forces exerted against the thumb, facilitating the interpretation of digit coordination in grasp control. The grip force of the virtual finger is the sum of the normal forces exerted by the individual fingers, and the load force is the sum of their tangential forces.

### Statistical analysis

All statistical analyses were carried out using repeated-measures analyses of variance (ANOVAs), with the significance threshold set at an alpha level of 0.05. The analyses were implemented in R [44]. To examine grip forces vis-à-vis our central hypothesis on MAH, a two-way repeated-measures ANOVA was conducted with Finger (Ring, Little) and Condition (ST, DTL, DTH) specified as within-subject factors. This design allowed for testing the main effects of Finger and Condition, as well as their interaction. The assumption of sphericity was assessed using Mauchly’s test, and when violations occurred, Huynh–Feldt corrections were applied. To further investigate grip force modulation at critical task epochs, four separate one-way repeated-measures ANOVAs were conducted on the virtual finger grip force (VF_GF) measured at distinct time points: lift onset, lift off, maximum VF_GF, and during the central hold phase. These ANOVAs tested the effect of Condition on VF_GF at each epoch, shedding light on the temporal dynamics of grip control under varying cognitive loads. In cases where significant effects were detected, post-hoc pairwise comparisons were conducted with Tukey’s HSD to account for multiple comparisons. For non-significant contrasts, equivalence tests were performed using pre-defined equivalence bounds based on the smallest effect size of interest (SESOI). Effect sizes were reported as partial eta squared (η_p_^2^) to indicate the magnitude of observed effects.

## Acknowledgments

We thank the Department of Science & Technology, Government of India, for supporting this work, vide Reference nos. SR/CSRI/97/2014 & DST/CSRI/2017/87 under Cognitive Science Research Initiative (CSRI) (awarded to Varadhan SKM). The funders had no role in study design, data collection and analysis, decision to publish, or preparation of the manuscript.

## Data Availability

The data will be made available in due course of time through a data descriptor article. Meanwhile, the data supporting the analyses presented in the paper are available upon reasonable request.

## References

[1] T. C. Pataky, M. L. Latash, and V. M. Zatsiorsky, “Tangential load sharing among fingers during prehension,” Ergonomics, vol. 47, no. 8, pp. 876–889, Jun. 2004, doi: 10.1080/00140130410001670381.

[2] F. Gao, M. L. Latash, and V. M. Zatsiorsky, “Internal forces during object manipulation,” Exp. Brain Res., vol. 165, no. 1, pp. 69–83, Aug. 2005, doi: 10.1007/s00221-005-2282-1.

[3] R. S. Johansson and G. Westling, “Coordinated isometric muscle commands adequately and erroneously programmed for the weight during lifting task with precision grip,” Exp. Brain Res., vol. 71, no. 1, Jun. 1988, doi: 10.1007/BF00247522.

[4] C. J. Winstein, J. H. Abbs, and D. Petashnick, “Influences of object weight and instruction on grip force adjustments,” Exp. Brain Res., vol. 87, no. 2, Nov. 1991, doi: 10.1007/BF00231864.

[5] M. Santello and J. F. Soechting, “Force synergies for multifingered grasping,” Exp. Brain Res., vol. 133, no. 4, pp. 457–467, Aug. 2000, doi: 10.1007/s002210000420.

[6] T. R. Schneider, G. Buckingham, and J. Hermsdörfer, “Torque-planning errors affect the perception of object properties and sensorimotor memories during object manipulation in uncertain grasp situations,” J. Neurophysiol., vol. 121, no. 4, pp. 1289–1299, Apr. 2019, doi: 10.1152/jn.00710.2018.

[7] Z. V., G. F., and L. M., “Prehension synergies: Effects of object geometry and prescribed torques,” Exp. Brain Res., vol. 148, no. 1, pp. 77–87, Jan. 2003, doi: 10.1007/s00221-002-1278-3.

[8] V. M. Zatsiorsky, F. Gao, and M. L. Latash, “Finger force vectors in multi-finger prehension,” J. Biomech., vol. 36, no. 11, pp. 1745–1749, Nov. 2003, doi: 10.1016/S0021-9290(03)00062-9.

[9] B. B. Edin, G. Westling, and R. S. Johansson, “Independent control of human finger-tip forces at individual digits during precision lifting.,” J. Physiol., vol. 450, no. 1, pp. 547–564, May 1992, doi: 10.1113/jphysiol.1992.sp019142.

[10] J. R. Flanagan, A. M. Wing, S. Allison, and A. Spenceley, “Effects of surface texture on weight perception when lifting objects with a precision grip,” Percept. Psychophys., vol. 57, no. 3, pp. 282–290, Apr. 1995, doi: 10.3758/BF03213054.

[11] R. S. Johansson and G. Westling, “Roles of glabrous skin receptors and sensorimotor memory in automatic control of precision grip when lifting rougher or more slippery objects,” Exp. Brain Res., vol. 56, no. 3, Oct. 1984, doi: 10.1007/BF00237997.

[12] V. M. Zatsiorsky, F. Gao, and M. L. Latash, “Prehension Stability: Experiments With Expanding and Contracting Handle,” J. Neurophysiol., vol. 95, no. 4, pp. 2513–2529, Apr. 2006, doi: 10.1152/jn.00839.2005.

[13] G. P. Slota, M. L. Latash, and V. M. Zatsiorsky, “Tangential Finger Forces Use Mechanical Advantage During Static Grasping,” J. Appl. Biomech., vol. 28, no. 1, pp. 78–84, Feb. 2012, doi: 10.1123/jab.28.1.78.

[14] V. M. Zatsiorsky, R. W. Gregory, and M. L. Latash, “Force and torque production in static multifinger prehension: biomechanics and control. II. Control,” Biol. Cybern., vol. 87, no. 1, pp. 40–49, Jul. 2002, doi: 10.1007/s00422-002-0320-7.

[15] P. Shenoy and V. S. K. M., “Task demands modulate distal limb handedness: A comparative analysis of prehensile synergies of the dominant and non-dominant hand,” Sci. Rep., vol. 14, no. 1, p. 25565, Oct. 2024, doi: 10.1038/s41598-024-75001-3.

[16] R. Banuvathy and S. Varadhan, “Distinct behavior of the little finger during the vertical translation of an unsteady thumb platform while grasping,” Sci. Rep., vol. 11, no. 1, p. 21064, Oct. 2021, doi: 10.1038/s41598-021-00420-5.

[17] B. L. Prilutsky, “Coordination of Two- and One-Joint Muscles: Functional Consequences and implications for Motor Control,” Motor Control, vol. 4, no. 1, pp. 1–44, Jan. 2000, doi: 10.1123/mcj.4.1.1.

[18] T. S. Buchanan, G. P. Rovai, and W. Z. Rymer, “Strategies for muscle activation during isometric torque generation at the human elbow,” J. Neurophysiol., vol. 62, no. 6, pp. 1201–1212, Dec. 1989, doi: 10.1152/jn.1989.62.6.1201.

[19] W. Zhang, H. B. Olafsdottir, V. M. Zatsiorsky, and M. L. Latash, “Mechanical Analysis and Hierarchies of Multidigit Synergies during Accurate Object Rotation,” Motor Control, vol. 13, no. 3, pp. 251–279, Jul. 2009, doi: 10.1123/mcj.13.3.251.

[20] H. Olafsdottir, W. Zhang, V. M. Zatsiorsky, and M. L. Latash, “Age-related changes in multifinger synergies in accurate moment of force production tasks,” J. Appl. Physiol., vol. 102, no. 4, pp. 1490–1501, Apr. 2007, doi: 10.1152/japplphysiol.00966.2006.

[21] J. K. Shim, M. L. Latash, and V. M. Zatsiorsky, “Finger coordination during moment production on a mechanically fixed object,” Exp. Brain Res., vol. 157, no. 4, pp. 457–467, Aug. 2004, doi: 10.1007/s00221-004-1859-4.

[22] B. Rajakumar, S. Dutta, and S. K. M. Varadhan, “Support for mechanical advantage hypothesis of grasping cannot be explained only by task mechanics,” Sci. Rep., vol. 12, no. 1, p. 10242, Jun. 2022, doi: 10.1038/s41598-022-14014-2.

[23] Y. Li, J. Randerath, H. Bauer, C. Marquardt, G. Goldenberg, and J. Hermsdörfer, “Object properties and cognitive load in the formation of associative memory during precision lifting,” Behav. Brain Res., vol. 196, no. 1, pp. 123–130, Jan. 2009, doi: 10.1016/j.bbr.2008.07.031.

[24] B. Lee, R. Miyanjo, F. Tozato, and Y. Shiihara, “Dual-task Interference in a Grip and Lift Task,” KITAKANTO Med. J., vol. 64, no. 4, pp. 309–312, 2014, doi: 10.2974/kmj.64.309.

[25] M. A. Spiegel, D. Koester, and T. Schack, “The functional role of working memory in the (re-)planning and execution of grasping movements.,” J. Exp. Psychol. Hum. Percept. Perform., vol. 39, no. 5, pp. 1326–1339, 2013, doi: 10.1037/a0031398.

[26] M. A. Spiegel, D. Koester, and T. Schack, “Movement planning and attentional control of visuospatial working memory: evidence from a grasp-to-place task,” Psychol. Res., vol. 78, no. 4, pp. 494–505, Jul. 2014, doi: 10.1007/s00426-013-0499-3.

[27] R. Gunduz Can, T. Schack, and D. Koester, “Movement Interferes with Visuospatial Working Memory during the Encoding: An ERP Study,” Front. Psychol., vol. 8, p. 871, May 2017, doi: 10.3389/fpsyg.2017.00871.

[28] J.-P. Van Dijck, W. Fias, and M. Andres, “Selective interference of grasp and space representations with number magnitude and serial order processing,” Psychon. Bull. Rev., vol. 22, no. 5, pp. 1370–1376, Oct. 2015, doi: 10.3758/s13423-014-0788-x.

[29] C. Voelcker-Rehage, A. J. Stronge, and J. L. Alberts, “Age-related Differences in Working Memory and Force Control under Dual-task Conditions,” Aging Neuropsychol. Cogn., vol. 13, no. 3–4, pp. 366–384, Dec. 2006, doi: 10.1080/138255890969339.

[30] A. K. Au and P. J. Keir, “Interfering effects of multitasking on muscle activity in the upper extremity,” J. Electromyogr. Kinesiol., vol. 17, no. 5, pp. 578–586, Oct. 2007, doi: 10.1016/j.jelekin.2006.06.005.

[31] R. K. Mehta and M. J. Agnew, “Effects of concurrent physical and mental demands for a short duration static task,” Int. J. Ind. Ergon., vol. 41, no. 5, pp. 488–493, Sep. 2011, doi: 10.1016/j.ergon.2011.04.005.

[32] J.-J. Temprado, S. Vieluf, N. Bricot, E. Berton, and R. Sleimen-Malkoun, “Performing Isometric Force Control in Combination with a Cognitive Task: A Multidimensional Assessment,” PLOS ONE, vol. 10, no. 11, p. e0142627, Nov. 2015, doi: 10.1371/journal.pone.0142627.

[33] D. J. Serrien, “Verbal–manual interactions during dual task performance: An EEG study,” Neuropsychologia, vol. 47, no. 1, pp. 139–144, Jan. 2009, doi: 10.1016/j.neuropsychologia.2008.08.004.

[34] S. A. Fraser, K. Z. H. Li, and V. B. Penhune, “Dual-Task Performance Reveals Increased Involvement of Executive Control in Fine Motor Sequencing in Healthy Aging,” J. Gerontol. B. Psychol. Sci. Soc. Sci., vol. 65B, no. 5, pp. 526–535, Sep. 2010, doi: 10.1093/geronb/gbq036.

[35] Y. Korotkevich, K. M. Trewartha, V. B. Penhune, and K. Z. H. Li, “Effects of age and cognitive load on response reprogramming,” Exp. Brain Res., vol. 233, no. 3, pp. 937–946, Mar. 2015, doi: 10.1007/s00221-014-4169-5.

[36] S. Dutta and V. Skm, “Working memory load disrupts motor performance but preserves adaptation during dexterous object manipulation,” Sep. 09, 2025, In Review. doi: 10.21203/rs.3.rs-7420188/v1.

[37] E. Guillery, A. Mouraux, and J.-L. Thonnard, “Cognitive-Motor Interference While Grasping, Lifting and Holding Objects,” PLoS ONE, vol. 8, no. 11, p. e80125, Nov. 2013, doi: 10.1371/journal.pone.0080125.

[38] E. Guillery, A. Mouraux, J.-L. Thonnard, and V. Legrain, “Mind Your Grip: Even Usual Dexterous Manipulation Requires High Level Cognition,” Front. Behav. Neurosci., vol. 11, p. 220, Nov. 2017, doi: 10.3389/fnbeh.2017.00220.

[39] B. Rajakumar and V. Skm, “Comparable behaviour of ring and little fingers due to an artificial reduction in thumb contribution to hold objects,” PeerJ, vol. 8, p. e9962, Sep. 2020, doi: 10.7717/peerj.9962.

[40] B. Rajakumar and S. K. M. Varadhan, “Evidence to support the mechanical advantage hypothesis of grasping at low force levels,” Sci. Rep., vol. 12, no. 1, p. 20834, Dec. 2022, doi: 10.1038/s41598-022-25351-7.

[41] R. C. Oldfield, “The assessment and analysis of handedness: The Edinburgh inventory,” Neuropsychologia, vol. 9, no. 1, pp. 97–113, Mar. 1971, doi: 10.1016/0028-3932(71)90067-4.

[42] M. F. Folstein, S. E. Folstein, and P. R. McHugh, “‘Mini-mental state,’” J. Psychiatr. Res., vol. 12, no. 3, pp. 189–198, Nov. 1975, doi: 10.1016/0022-3956(75)90026-6.

[43] C. L. MacKenzie and T. Iberall, The grasping hand. in The grasping hand. Amsterdam, Netherlands: North-Holland/Elsevier Science Publishers, 1994, pp. xvii, 482.

[44] R Core Team, R: A language and environment for statistical computing. Vienna, Austria: R Foundation for Statistical Computing, 2021. [Online]. Available: https://www.R-project.org/

